# Generalizing clusters of similar species as a signature of coexistence under competition

**DOI:** 10.1101/264606

**Authors:** Rafael D’Andrea, Maria Riolo, Annette Ostling

## Abstract

Patterns of trait distribution among competing species can potentially reveal the processes that allow them to coexist. It has been recently proposed that competition may drive the spontaneous emergence of niches comprising clusters of similar species, in contrast with the dominant paradigm of greater-than-chance species differences. However, current clustering theory relies largely on heuristic rather than mechanistic models. Furthermore, studies of models incorporating demographic stochasticity and immigration, two key players in community assembly, did not observe clusters. Here we demonstrate clustering under partitioning of resources, partitioning of environmental gradients, and a competition-colonization tradeoff. We show that clusters are robust to demographic stochasticity, and can persist under immigration. While immigration may sustain clusters that are otherwise transient, too much dilutes the pattern. In order to detect and quantify clusters in nature, we introduce and validate metrics which have no free parameters nor require arbitrary trait binning, and weigh species by their abundances rather than relying on a presence-absence count. By generalizing beyond the circumstances where clusters have been observed, our study contributes to establishing them as an update to classical trait patterning theory.

**Author Summary:** Species traits determine how they compete with each other. As such, patterns in the distributions of traits in a community of competing species may reveal the processes responsible for coexistence. One central idea in theoretical ecology is that the strength of competition relates to similarity in species needs and strategies, and therefore if competition plays out at short timescales, coexisting species should be more different than expected by chance. However, recent theory suggests that competition may lead species to temporarily self-organize into groups with similar traits. Here we show that this clustering is a generic feature of competitive dynamics, which is robust to demographic stochasticity and can be indefinitely maintained by immigration. We show that clustering arises whether species coexist by partitioning resources, environmental preferences, or through tradeoffs in life-history strategies. We introduce and validate metrics that, given species traits and abundances, determine whether they are clustered, and if so, how many clusters occur. By showing the generality of self-organized species clusters and providing a tools for their detection, our study contributes to updating classical ideas about how competition shapes communities, and motivates searches for them in nature.

## Introduction

Competition is a driving force in nature, and a central question in ecology is how it shapes community structure. Species traits mediate their interactions with the environment and each other, and therefore determine how they compete. As such, patterns of trait distribution among co-occurring species can give insights into the underlying coexistence mechanisms [1–5]. Theory shows that coexistence is only stable if species differ in their ecological needs and impacts—i.e. they must display niche differences (see Glossary) [6,7]. These differences reduce competition between species compared to competition within species, thus allowing each species to grow from low abundance in the presence of the others. If species are arranged on niche axes so that proximity on those axes indicates similarity in their niche strategies, they are expected to display limiting similarity–greater-than-chance differences on those axes [8]. Ecologists use trait axes as proxies for niche axes, and look for similar patterns therein, usually in the form of overdispersion or even spacing between species on these axes [9–17]. This classical idea remains the dominant paradigm, despite mixed empirical support [2,3,5].

In contrast, recent studies suggest that competition causes the spontaneous emergence of transient clusters of species with similar traits [18–20]. They find that limiting similarity results only if competition plays out to the final stage, whereby all species are excluded except for those with optimal niche strategies (see Glossary). Those species are niche-differentiated enough from one another (i.e. have enough space between them on the niche axis) to all stably coexist together, having emerged as dominant over other strategies through the competitive process. When species outnumber optimal niches (e.g. phytoplankton communities with more species than resources, 21), competitive exclusion will ensue. However, species near the optimal niche strategies are excluded more slowly than those further away from these optimal strategies [22–24]. As a result, the community temporarily self-organizes into clusters of similar species near optimal strategies, with gaps in between.

Clustering has traditionally been associated with environmental filters [25], and more recently with one-sided competition without a balancing tradeoff, such as competition for light where taller is better [26]. However, those will tend to produce a single big cluster around the favored trait, whereas partitioning of a niche axis will lead to multiple clusters, one per optimal niche strategy. Patterns of multiple clusters have in fact been reported in empirical studies of body size in phytoplankton communities and certain animal taxa [18,27–29]. This phenomenon has been interpreted to bridge coexistence through differences (niche theory) and similarity (neutral theory) [30], and as such potentially represents a unification of classical ideas and a generalization of limiting similarity.

Scheffer and van Nes [18] first demonstrated the emergence of transient clusters in Lotka-Volterra dynamics. Later work further suggested that clusters are a generic outcome of Lotka-Volterra dynamics [20] except for special shapes of the function connecting competition to traits [31,32]. Several mechanisms can make these transient clusters persist, such as specialist enemies [18] and periodic environments [33,34].

However, the generality of clusters as a signature of competition cannot be established without showing that they emerge in communities subject to stochastic processes. In particular, immigration and ecological drift are intrinsic to most communities, and have been amply demonstrated to be important players in nature [30,35–37]. In fact, models ignoring all of biology but drift and immigration successfully describe observed macroecological patterns [38] (which of course does not mean deterministic forces are unimportant). Yet, clustering remains unseen in competition models incorporating these processes [39–41]. In plankton models where clustering occurs, immigration has as a negative impact [21].

Moreover, clusters have not been widely demonstrated beyond Lotka-Volterra dynamics. Lotka-Volterra competition equations are a heuristic description which does not specify a niche mechanism. They are a special limit [42] of MacArthur’s consumer-resource dynamics model [43], and cannot describe all types of competition [44]. While species clusters have been shown to emerge in explicit consumer-resource dynamics [21,34], these studies ignored the possibility of resource depletion effects. The latter have been shown to greatly affect competitive relations between consumers, for example by violating the assumption that competition always decreases with trait differences [45]. It is also not clear that clustering should emerge among species competing along environmental gradients; indeed, early studies of stochastic niche dynamics [39–41] focused on competition of this type and found no clustering pattern (although these studies, which predate [18], were not specifically looking for clustering). Finally, it is not known whether clusters emerge in communities characterized by hierarchical competition among species that coexist via life-history tradeoffs. These mechanisms, such as the competition-colonization tradeoff [46] and the tolerance-fecundity tradeoff [47], have been chiefly studied in terrestrial plants, but may enable coexistence among other sessile organisms with a dispersive stage, such as coral, coral fishes, and microbes [47].

To determine whether clusters are a general outcome of competition as opposed to an artifact of specific models, and to verify their robustness to stochastic forces, here we use a stochastic niche simulation approach to investigate the emergence of clustering by traits in species assemblages undergoing competitive dynamics and open to immigration.

We start with Lotka-Volterra dynamics, where clusters are known to emerge in the deterministic model [18], to address the question of their robustness to stochastic forces. We test this robustness with and without the confounding influence of environmental filters, which we hypothesize might mask any clustering caused by niche partitioning. We then see how regional diversity and immigration rate influence this robustness. Clustering intrinsically involves many species, and should in principle be stronger, or at least more detectable, under a high species-to-niches ratio. Therefore, we expect more clustering under higher regional diversity. As for immigration, we expect that while it may contribute to the persistence of clusters by keeping weak competitors from being excluded, too much immigration will drown any pattern caused by competition.

Next, to determine whether clusters are a general outcome of competition, we analyze models spanning three key niche mechanisms: resource partitioning, habitat partitioning, and a competition-colonization tradeoff. Under resource partitioning, we further explore the differences between scenarios with low and high resource depletion. Under competition for habitat, we also examine the impact of dispersal limitation as opposed to global dispersal. We expect that larger scale dispersal relative to the scale of the environmental gradient can influence cluster emergence by lessening the dominance of emergent optimal strategies, since their propagules often spread to locations with suboptimal habitat. If clusters are a signature of competition and niche differentiation, we expect clustering to emerge generally under all scenarios of all three niche mechanisms.

To determine whether communities are clustered beyond patterns that could arise by chance, we present two metrics: the first uses the k-means algorithm [48], which assigns species to clusters by minimizing trait differences within clusters. The second is based on Ripley’s K [49], and quantifies clustering based on the sparseness of the regions between clusters. Both metrics apply the gap statistic method for comparing data with null models [50]. These metrics improve over existing tools to detect clustering [18, 27–29, 51] as they contain no free parameters and do not arbitrarily bin traits. Furthermore, rather than reducing species to presence-absence counts, our metrics weigh them by their abundance. We validate the metrics by confirming that they detect clusters in Lotka-Volterra communities but not in neutral communities.

## Results

All niche mechanisms (Fig 1) resulted in species clustering (Fig 2A-G, Fig 3), demonstrating both the robustness to stochasticity and generality of clustering emergence under niche differentiation. In contrast, neutral communities (Fig 2H) were clustered only about as often as expected by chance, and communities under no niche mechanism but subject to environmental filters collapsed into a single cluster (Fig S1). Our success at detecting clusters under niche differentiation, combined with this lack of false detection when niche mechanisms are absent, suggests our metrics are well-suited to quantifying the pattern arising from niche differentiation.

**Figure 1:**
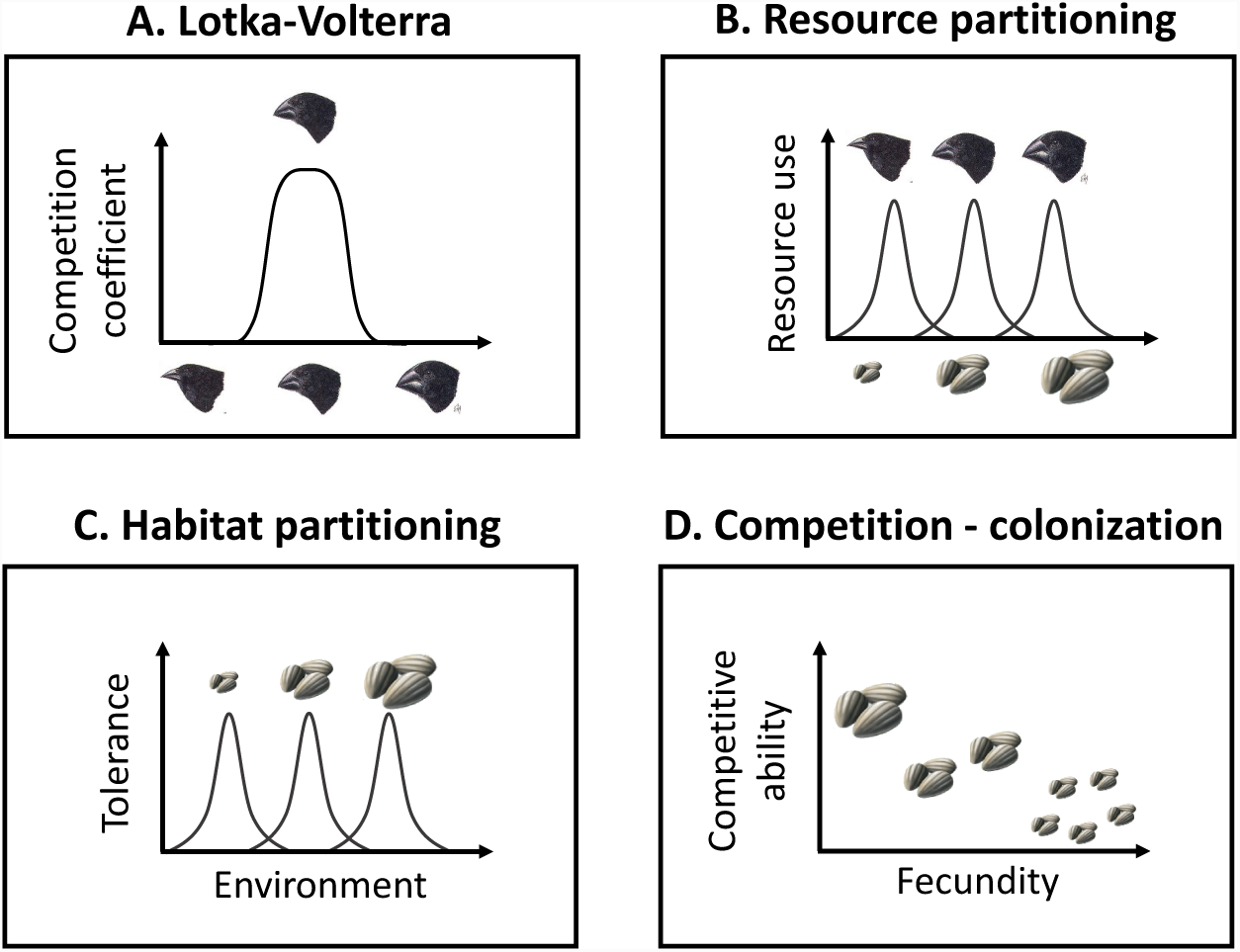
Schematic representation of niche models. Panel A illustrates Lotka-Volterra competition, while panels B-D illustrate different niche mechanisms, with the niche axis represented on the abscissa. **A**. Lotka-Volterra: competition coefficients are tied to traits, here beak size. Curve shows competitive impact by each species on the focal species, increasing with trait similarity. **B**. Partitioning resources: We assume a continuum of resources (seeds), and species traits (beak size) determine their use of the different resources. Depending on the degree of specialization, resource depletion may occur (see text). **C**. Partitioning environment: species with different traits (seed size) are optimally adapted to different environments. This model is spatially explicit, and we implement it both with and without dispersal limitation; **D**. Competition-colonization tradeoff: species trade off fecundity (seed output) with ability to win sites against other species (mediated by seed size).

**Figure 2:**
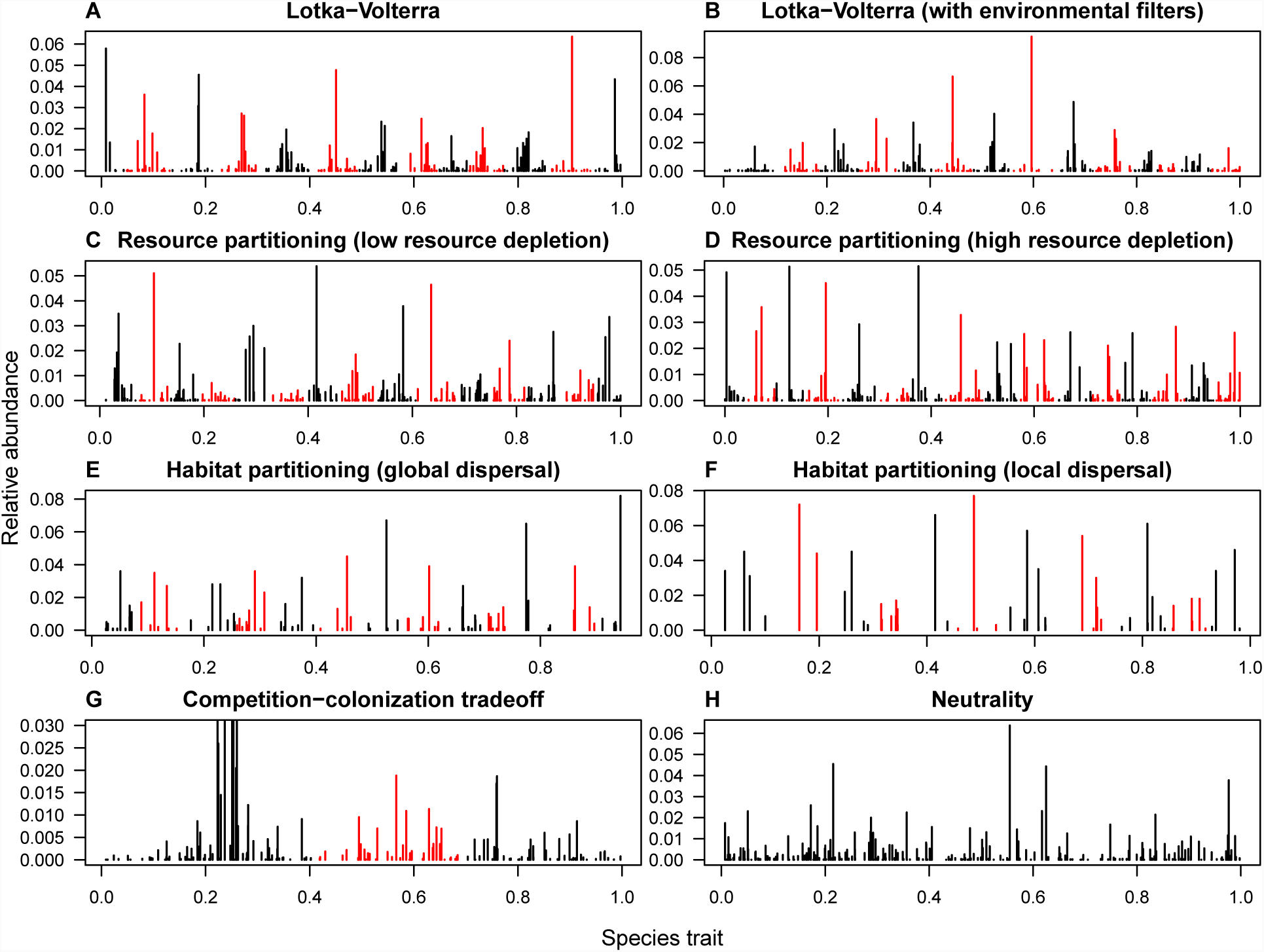
Abundance-by-trait pattern across niche mechanisms. For each scenario (A-H) we show one representative replicate. All communities with a niche mechanism (A-G) are clustered at the *p* < 0.05 level, while the neutral community (H) is not (*p* = 0.1). Alternating black and red colors highlight the clusters. We truncate the y-axis in the competition-colonization tradeoff scenario (G) to better show abundance structure among rare species. (Immigration rate *m* = 0.08, regional diversity c. 400 species. All communities had 21,000 individuals, except in the partitioning environment scenario, which had 1,000 individuals. Parameters of niche models were tuned so that without stochasticity and immigration they would produce about 13 transient clusters, and eventually 13 species stably coexisting at equilibrium.)

**Figure 3:**
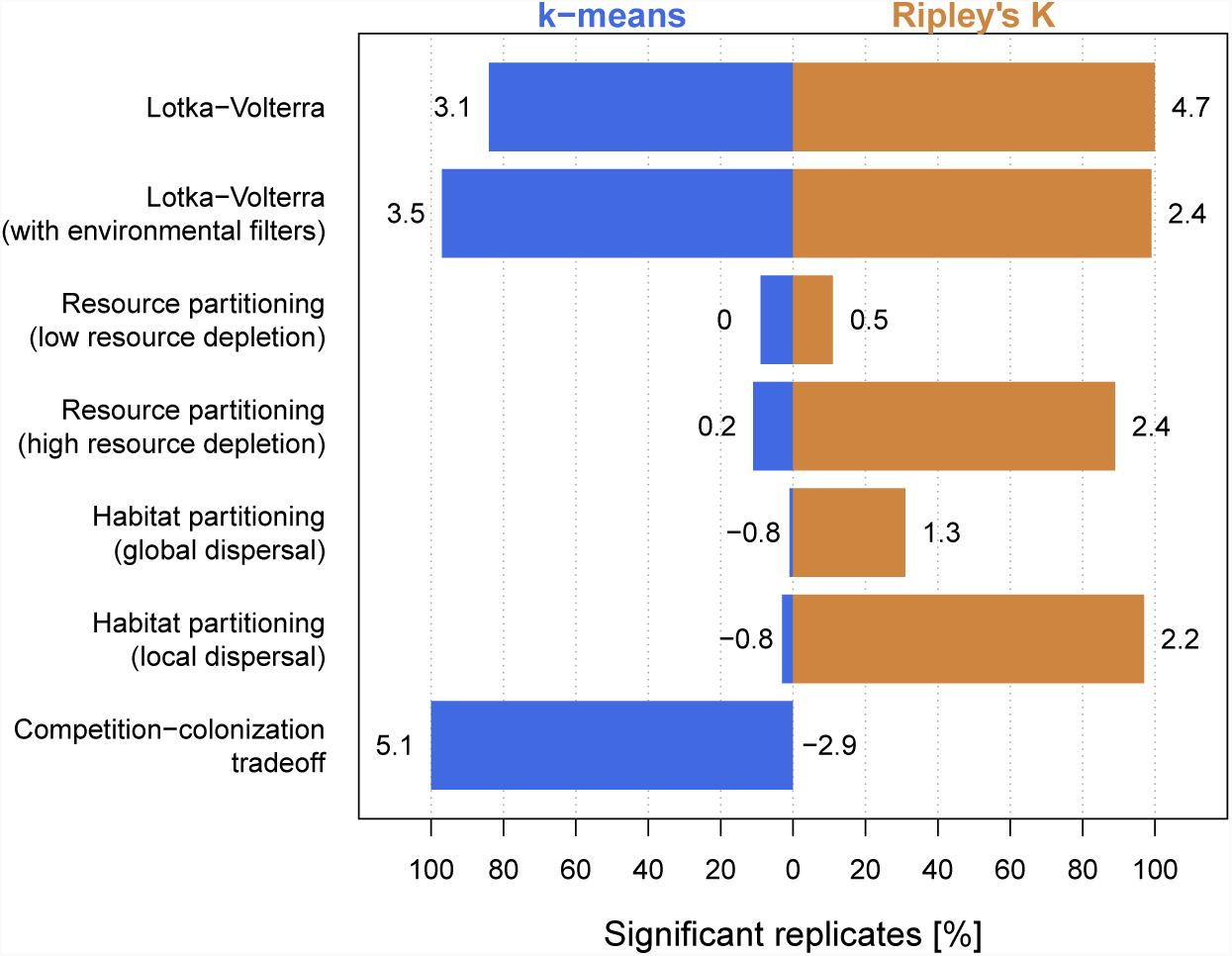
Clustering across niche mechanisms. Bars show percentage of replicates that were significantly clustered at level *p* < 0.05 according to the k-means metric (blue) and the Ripley’s K metric (orange), out of 100 replicates. Numbers next to bars show average z-score. For comparison, 8% of neutral communities were clustered by the k-means metric 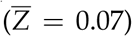, and 7% by the Ripley’s K metric 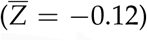, thus close to the 5% background detection expected from a null model (z-scores and p-values were obtained by comparing each replicate against 100 null communities. *m* = 0.08, c. 400 regional species across scenarios.)

### Lotka-Volterra

We implemented a stochastic analogue of classical Lotka-Volterra competition, which assumes species influence each other directly via competition coefficients. We tied those coefficients to similarity in traits (Fig 1A), so that trait differences can enable stable coexistence (niche differentiation). In its deterministic form, this type of model is known to produce clusters [18] (except for special cases where competition coefficients form a positive-definite matrix [32,52]). We thus used this model to consider the robustness of clusters to stochastic forces. We also used it to consider whether that robustness of clustering is influenced by the presence of environmental filtering, and by regional diversity and immigration.

Clustering was strong enough in the Lotka-Volterra communities to be easily distinguishable by eye (Fig 2A-B), with over 80% of all 100 replicates testing significant at the *p* < 0.05 level by either metric (Fig 3). We note that the transient clusters produced by deterministic Lotka-Volterra competition are being maintained in our stochastic model by immigration; when immigration is turned off, communities begin to lose species and tend towards a limiting similarity pattern [23].

### Environmental filtering

Environmental filtering [25] or physiological constraints may cause some species to grow faster than others in the absence of competitors. For example, species with intermediate traits may grow faster because extreme trait values may have metabolic costs or are maladapted to the local environment, regardless of competitive interactions. This will affect abundances, and might interfere with the emergence of niche-driven clusters. In our simulations, environmental filters against extreme traits caused species near the center of the trait axis to be more abundant than their counterparts near the edges, as expected (Fig 2B), but this had minimal impact on clustering relative to no filtering (compare average z-scores, Fig 3). However, as we increased the strength of environmental filters, they ultimately eclipsed niche partitioning as a driver of pattern, and the community appeared as a single cluster (Fig S2), consistent with results from environmental filters acting alone.

### Regional diversity

Clustering increased with regional diversity. As we increased the number of species in the regional pool while keeping the number of clusters fixed, the z-score of the clustering metric rose monotonically (Fig 4). This reflects the fact that clustered patterns in general are increasingly distinguishable from randomness as the number of items per cluster increases (Fig S3).

**Figure 4:**
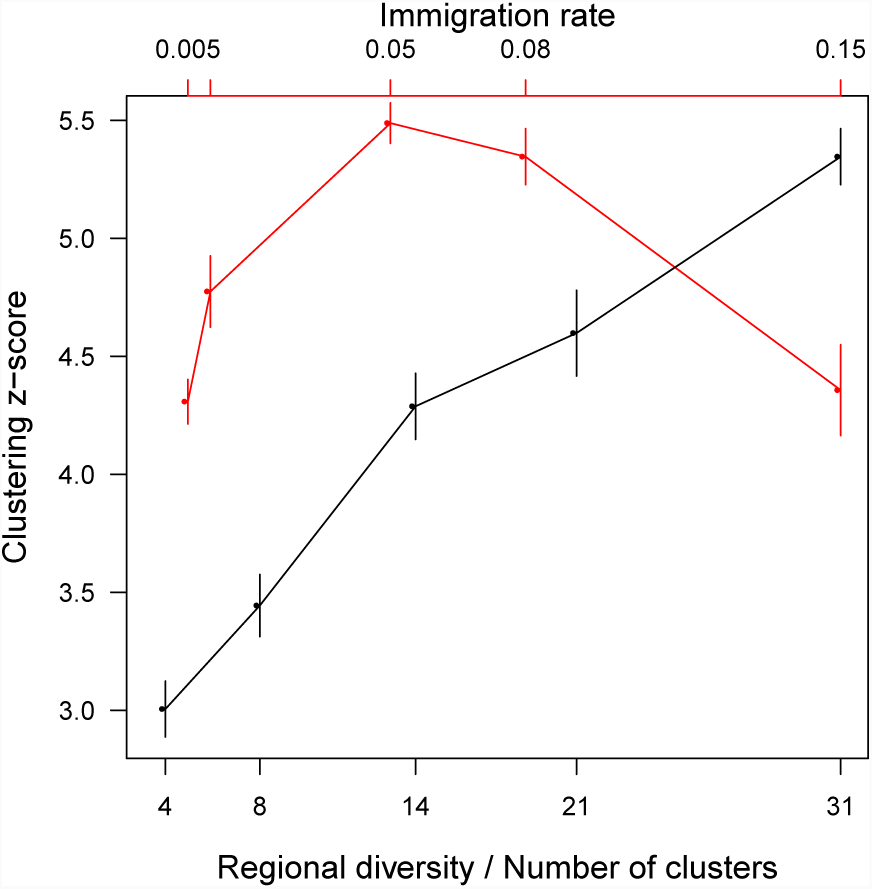
Impact of immigration and regional diversity. Clustering increases with regional diversity (black), and has a modal relationship with immigration pressure (red). Bottom axis: number of species in the pool divided by the number of clusters in the community. Top axis: immigration rate. Points show the mean z-score of the Ripley’s K metric across 10 replicates. Error bars show 1.96 standard errors of the mean. (*m* = 0.08 when varying regional diversity, and the latter is fixed at c. 400 regional species when varying immigration.)

### Immigration rates

Clustering had a modal relationship with immigration pressure: as we increased the immigration rate from low values, the clustering z-score first rose, then peaked and fell (Fig 4). This indicates that while immigration sustains the clusters by replenishing populations that would otherwise be excluded, too much immigration dilutes the effects of local dynamics and adds noise to the pattern (Fig S3). While this modal shape is arguably general, the position of the mode is specific to the niche mechanism and other factors such as environmental filters.

Communities in the other niche models under immigration rate *m* = 0.01 as opposed to 0.08 generally showed a similar number of clusters but fewer species per cluster, and gaps between clusters were more rarified (Fig S4). In some cases, this increased clustering (Fig S5), suggesting that *m* = 0.08 falls to the right of the mode in those niche models.

### Resource partitioning

Assuming fast resource dynamics relative to consumers, one can derive the Lotka-Volterra model from resource-consumer dynamics [42,53], with trait similarity indicating similar resource preferences [8,43] (Fig 1B). However, this approach ignores the effects of stochastic fluctuations in resource availability, and particularly resource depletion. The latter has been shown to impact co-existence outcomes [54] and even competition-similarity relationships [45]. These relationships are drivers of pattern [31], and we therefore tested whether incorporating stochastic resource dynamics and depletion affected the emergence of clusters.

In the case of low resource depletion, we found weak clustering (Fig 3). This is because when resource depletion is low, consumers undergo approximate Lotka-Volterra dynamics with competition coefficients following a Gaussian function of trait separation [8]. This function is positive-definite, a property that has been shown to lead to weak or no pattern (see Fig S8 in [23] and [32,52]). Species sorting under Gaussian competition is slow (see Fig S8 in [23]), and therefore easily overpowered by immigration. Indeed, clustering was stronger under lower immigration pressure (Figs. S4B, S5).

Resource depletion breaks the link to Gaussian Lotka-Volterra dynamics, thereby fundamentally changing the competitive interactions between consumers [45,54]. That strengthened clustering in our model (compare low- and high-depletion scenarios in Fig 3). As resources were extirpated, species ended up clustering based on specialization to the remaining ones (Fig S6). Resource depletion thus sets the optimal niche strategies (i.e. specialization onto remaining resources), thereby strengthening the pattern.

### Habitat partitioning

Habitat is a critical resource for which species compete, and as such the environment is thought of as a key axis of niche differentiation [55–57]. Traits reflect adaptations to different environments, and hence competition for habitat must shape trait distribution.

To test whether niche-differentiation based on environmental preference leads to clustering, we used the model introduced by [40]. We assume a linear landscape on a habitat gradient, e.g. an elevation gradient. Different species are optimally adapted to different habitats, and competition arises from overlap in environmental preference (Fig 1C). Competition occurs at the recruitment stage, where the probability of recruiting is based on tolerance to the local environment. Because dispersal can play a central role in competition for space [57,58], we consider two scenarios: global and local dispersal, which differ by whether individuals are more likely to disperse shorter distances from their parents. Since this is an individual-based model, we used a smaller community size (1,000 individuals rather than 21,000) for computational expedience.

We found that clusters also emerge under competition for habitat. Switching dispersal from global to local had a strong impact on cluster shape, and reduced the number of species per cluster (compare Fig 2E, F). This occurred as dispersal limitation effectively decreases immigration. Moreover, the attending reduction of the diluting effects of immigration substantially strengthened clustering (Fig 3).

### Competition-colonization tradeoff

All models examined so far describe symmetric competition, whereby the competitive impact of species A on B is similar to that of B on A. Competitive hierarchies stand in contrast to this. That is the case of the competition-colonization tradeoff [46,59], where propagule production, or colonization ability, trades off with the ability to displace individuals of other species, or competitive ability (Fig 1D). Even though in this case some species are better competitors than others, the tradeoff is a niche mechanism because it allows for the stable coexistence of multiple species^1^ (see Glossary, Fig S7).

Clusters also emerged under this niche mechanism. The asymmetry in species interactions was reflected in its asymmetric clusters (Fig 2G). The k-means metric picked up on just three clusters in most replicates, even though without stochasticity and immigration the model produces about 13 transient clusters (Fig S7). This is perhaps because species in the first cluster so strongly dominated the community (e.g., the most abundant species in the first cluster in Fig 2G had 4,798 individuals, compared with 395 in the second and 392 in the third). This indicates that species adopting the high-competitiveness strategy (left side of the trait axis) outperform both those who invest in high fecundity and the intermediate group. Dispersal limitation could reduce the asymmetry by augmenting the benefits of high fecundity. We hypothesize that a spatially explicit formulation of this mechanism would produce more similarly sized clusters.

### Metric performance

While our metrics performed equally well in the Lotka-Volterra communities, they differed in the other niche mechanisms. Our Ripley’s K metric fared better than the k-means metric in scenarios where species partition resources and habitat (Fig 3) (though performance was similar for resource partitioning under low immigration, see Fig S5). The Ripley’s K metric focuses on identifying a scale of interspecific trait difference with particularly low representation (i.e. by which very few species pairs are separated). As such, it relies on regular spacing between clusters and intercluster gaps. Ripley’s K is good at identifying clusters when that spacing is regular, even in cases where the overall pattern is noisy (the resource partitioning cases), or involves low-occupancy clusters (the habitat partitioning cases). On the other hand, the k-means metric found the clusters in the competition-colonization tradeoff while Ripley’s K missed them (Fig 3). This is because the k-means algorithm is less sensitive to strong asymmetries between the clusters. In these cases, the k-means metric is a better choice.

## Discussion

Ecologists have long sought to understand how competition shapes community structure. While competing species are usually expected to be more different than predicted by chance [25,60], recent studies suggest that competition may cause species to cluster by traits, such that the community self-organizes into groups of similar species [18], a phenomenon which has been interpreted to bridge coexistence through differences–niche theory–and similarity–neutral theory. Our study verified that clustering transcends Lotka-Volterra dynamics, occurring under a number of niche mechanisms. Further, we showed that clustering is robust to stochastic drift, an intrinsic property of real-life communities. Immigration maintains clusters that are otherwise transient, and the strength of clustering has a modal relationship with immigration pressure. We showed that clustering may be detectable under the confounding influence of environmental filters, and is enhanced by regional diversity. Finally, we provided metrics for detecting and quantifying clusters in nature.

Why do clusters arise? Different niche mechanisms share the common property that competition is stronger between species with more similar strategies. It thus seems paradoxical that clusters should emerge. However, it is precisely because species with similar niches compete more strongly that clusters appear [61]. While similar pairs compete more strongly, they experience similar competitive pressure (or relief) from the rest of the community. If a given niche strategy is favored because it minimizes competition with the rest of the community or capitalizes on greater resource supplies, then similar strategies are similarly favored. This hilly fitness landscape causes exclusion to be slower near the center of the niches than in the gaps between them, making it easy for immigration to permanently maintain the clusters [23].

Modern coexistence theory [7] splits coexistence-promoting processes into those that reduce competition among species relative to competition within species (stabilizing mechanisms, here referred to as niche mechanisms), and those that reduce differences in average fitness between species (equalizing mechanisms). It is thus tempting to interpret clusters as reflecting a harmonious combination of stabilizing and equalizing forces: species within a cluster are equalized, while those in different clusters are stabilized. However, this interpretation is problematic. The equalization-stabilization dichotomy is based on applying invasibility criteria to closed communities regulated by a small number of limiting factors [62], an approach which does not extend easily to multispecies communities under immigration and a continuum of resources. In diverse communities with complex competitive interactions, it is difficult to calculate equalizing and stabilizing terms and tie them to trait differences, let alone interpret how specific patterns such as clustering connect with equalization and stabilization forces [62]. One approach is to assume all pairs of species compete with equal intensity [7,63], but this assumption is strongly violated in all models where clustering has been observed so far.

While competition is responsible for clusters’ emergence, immigration is responsible for their persistence. Immigration joins other mechanisms that have been previously shown to sustain clusters, namely specialist enemies [18,64] and environmental fluctuations [33,34]. We found that clustering appears generally under different immigration regimes, especially if the number of species far exceeds the number of available niches. Studies of stochastic competitive preceding Scheffer and van Nes 2006 found little impact of immigration on resulting trait pattern [39], [40]. However, successful immigration was highly infrequent in those models. In [39], resources made available through deaths were assumed to be redistributed broadly, so that resource supply remained low everywhere, making recruitment of new individuals highly unlikely. In [40], immigration took the form of a single immigrant seed being added to a large pool of local seeds competing for the site. As such, immigration rates were effectively much lower than ours. [41] tested both very high and very low immigration, and also saw no clusters (in no small part because the authors were not looking for them!). This could be due to their use of a Gaussian competition kernel (i.e. competition coefficients are a Gaussian function of trait difference), which leads to weak niche sorting dynamics, easily overwhelmed under high immigration [32].

The fact that immigration maintains clusters seemingly defers the question of coexistence to the regional scale. The problem dissipates by considering mass effects in the metacommunity framework [65]. The regional pool is a combination of local communities, and species are selected for different traits at different sites due to their own local niche dynamics. Therefore, each community receives immigrants which may be dominant elsewhere despite being disfavored locally. However, this does not mean that pattern is expected regionally, as the sum total of heterogeneous communities, each with a different trait pattern, may result in no discernible pattern at a regional scale.

Clusters caused by partitioning of a niche axis are often distinct from clustering due to other processes. Environmental filters favoring a single best trait and one-sided competitive dynamics where a particular trait outperforms all others without a balancing tradeoff [26] will produce a single cluster as opposed to multiple clusters. Where evolutionary rates are commensurate with ecological dynamics, small mutations and sympatric speciation may also generate species or genotype clusters without niche differentiation [66,67]. Ruling out these alternative sources of clustering could require sampling at a larger scale than applicable to the niche mechanism [5,60], or directly verifying competitive effects and frequency dependence [4]. While clusters may have more than one source, observation of multiple clusters in specific functional traits can help identify potential drivers of niche differentiation in a community.

We presented and validated two nonparametric abundance-weighted tools for detecting and quantifying clusters. Our metrics successfully distinguished clustering in niche-differentiated communities from no clustering in neutral communities and a single cluster in communities under environmental filters without a niche mechanism. We note that none of these results would appear without considering species abundances, as presence-absence counts do not reveal clustering in our communities. Also, our metrics did not require arbitrarily binning traits, nor fitting parameters to the data.

Although our study focused on one-dimensional trait axes, competitive interactions may often be mediated by variation in multiple traits [68]. Theoretical work on simple models indicates that multidimensional niche space leads to multidimensional clustering [20]. Our metrics can quantify clusters in any dimension, using generalized measures of niche separation. The main challenge is to connect multidimensional phenotypes to competitive relations, in order to define the correct measure of distance in high-dimensional trait space [61]. However, even if species cannot be arranged on linear trait axes, our Ripley’s K metric can still detect clustering by similarity as long as a measure of species differences can be defined, e.g. Hamming distance in genetic sequences [66].

Clustering as a signature of coexistence under competition is an update to the still dominant paradigm that competing species will display greater-than-chance differences [3–5, 16, 17]. Our finding that clusters appear under various niche mechanisms and can be easily maintained by immigration, even when confounding forces are at play, suggests that clusters are a likely feature of nature beyond the instances where it is currently known to occur [18,28,29]. For example in tropical forests, where both competition and dispersal are recognized as key drivers of community assembly [69,70], clustering could help explain the high diversity and seemingly continuous phenotypic variation.

## Methods

### Models and simulation design

We used a lottery model framework [71] to implement stochastic niche dynamics in a fixed-size community open to immigration from a regional pool. We start with a random draw of offspring from the regional pool, and then alternate death and recruitment events until species abundance distributions are stationary. A proportion *m* of deaths are replaced by immigrants, and the remainder by local offspring. This is analogous to Hubbell’s neutral model [38], except here the niche mechanism sets the probabilities of birth and death across different species. Schematic illustrations of our niche models are shown in Fig 1.

We used the 50-hectare plot of tropical forest on Barro Colorado Island, Panama, as a reference point for our community size (21,000 individuals *>* 10 cm dbh [38]) and immigration rate (0.08 immigrant recruits per recruitment event [72]). Our regional pool is a fixed neutral meta-community with biodiversity parameter *θ* = 50 [38] and 150,000 individuals, leading to c. 400 species (bigger metacommunities did not change results).

All simulations and statistical analyses were done in R [73].

### Lotka-Volterra

The probability that species *i* is selected for a death event is Σ_*j*_ *A*_*ij*_ *N*_*i*_ *N*_*j*_/ Σ_*k*_ Σ_*l*_ *A*_*kl*_ *N*_*k*_ *N*_*l*_, where *N*_*i*_ is the abundance of species *i* and the competition coefficient *A*_*ij*_ quantifies the competition caused on species *i* by species *j*. The sums are over all species present in the local community, and the denominator normalizes the probabilities to 1. The probability that species *i* is chosen at the subsequent recruitment stage is *m q*_*i*_ + (1 *- m*) *r*_*i*_ *N*_*i*_/ Σ_*j*_ *r*_*j*_ *N*_*j*_, where *q*_*i*_ is the relative abundance of species *i* in the regional pool, and *r*_*i*_ is its intrinsic growth rate. (We are placing competitive interactions on mortality, but results are unchanged when we place them on births instead.)

We link competition to traits by setting *A*_*ij*_ = exp[*-* (*|x*_*i*_ *- x*_*j*_*|*/*w*)^4^], where *x*_*i*_, *x*_*j*_ *∈* [0, 1] are the trait values of species *i* and *j*. Thus competition peaks at *x*_*i*_ = *x*_*j*_ (intraspecific competition), and declines with increasing trait difference (Fig 1A). This is a generic unimodal function commonly used in competition models [31]. Other unimodal shapes bring no qualitative changes (with a few exceptions, see [32]). The scaling parameter *w* sets the width of niche overlap, and therefore the number of niches available to species. We set *w* = 0.063, leading to c. 13 clusters. Where possible, we set parameters to keep the number of clusters consistent.

In the scenario with environmental filtering we set intrinsic growth rates *r*_*i*_ = *x*_*i*_(1 *- x*_*i*_), such that species with intermediate traits grow faster than those with extreme traits. In all other Lotka-Volterra scenarios, we set all *r*_*i*_ = 1 identical across species.

We set default values *m* = 0.08 for immigration rate, and *S* ⋍ 400 for regional diversity (via *θ* = 50 in the regional pool). When examining the impact of immigration pressure, we used *m* = 0.005, 0.01, 0.05, 0.08, 0.15. When examining regional diversity, we used *S* ⋍ 50, 100, 180, 250, 400 by setting *θ* = 5, 10, 20, 30, 50.

### Partitioning resources

Let *X*_*a*_ and *N*_*i*_ be the abundances of resource *a* and species *i*. We assume resource consumption affects species reproduction, and all species have equal mortality. As such, the probability that species *i* is chosen for a death event is simply *N*_*i*_/ Σ_*j*_ *N*_*j*_, and the probability that it is chosen for a recruitment event is *m q*_*i*_ + (1 - *m*) Σ_*a*_ *C*_*ai*_ *X*_*a*_ *N*_*i*_/ Σ_*j*_ Σ_*b*_ *C*_*bj*_*X*_*b*_ *N*_*j*_, where *C*_*ai*_ quantifies the preference of species *i* for resource *a*.

Resources simultaneously follow their own stochastic birth-death process. Consumption lowers resource populations, which would otherwise grow logistically. Therefore, resource *a* is selected for a death event with probability Σ_*i*_ *C*_*ai*_ *X*_*a*_ *N*_*i*_/ Σ_*b*_ Σ_*j*_ *C*_*bj*_*X*_*b*_ *N*_*j*_, and for a birth event with probability (1 - *X*_*a*_/*K*)*X*_*a*_/ Σ_*b*_(1 − *X*_*b*_/*K*)*X*_*b*_, where *K* is the resource carrying capacity, set equal across all resources for simplicity. There is no resource immigration; if a resource population is depleted to zero, the resource is extirpated and cannot be replenished.

Resources are arranged on a line between 0 and 1. We define *C*_*ai*_ = exp[*-* (*d*_*ai*_/*σ*)^2^], where *d*_*ai*_ is the difference between resource *a* and the preferred resource of consumer *i* (Fig 1B). The width parameter *σ* controls degree of resource overlap and thus number of niches. We set *σ* = 0.03 as it leads to about 13 clusters, for consistency with the Lokta-Volterra scenarios. We control resource depletion via the resource carrying capacity. When *K* = 100, most resources are never severely depleted by consumption, whereas when *K* = 400 resource depletion is severe and many resources are extirpated. Both scenarios start with 500 resources drawn from a standard uniform distribution.

### Partitioning habitat

We used the model introduced by [40]. We assume a linear array of sites forming a habitat gradient. Different species are optimally adapted to different environments, and competition arises from overlap in environmental preference (Fig 1C). Here, a death event is followed by a lottery in the local seed bank, which is formed via dispersal from all individuals in the community, plus a proportion of immigrants. A seed’s probability to win the lottery is given by its tolerance to the local environment, which declines with differences from the optimal environment of that species. All individuals have the same probability of dying. The probability that species *i* fills the vacancy is then 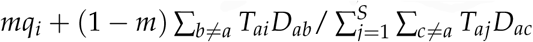, where *D*_*ab*_ is the probability of dispersal between an individual at site *b* and the vacated site *a*, and *T*_*ai*_ is species *i*’s tolerance to site *a*. We set *T*_*ai*_ = exp[*-* (Δ_*ai*_/*σ*)^2^], where Δ_*ai*_ is the difference between the local environment *a* and the niche optimum of species *i*. The width parameter *σ* controls niche overlap, and hence the number of niches. We set *σ* = 0.07 as it gives us c. 13 clusters.

We use a Gaussian dispersal kernel, *D*_*ab*_ = exp[*-* 0.5(*d*_*ab*_/*d*_disp_)^2^], where *d*_*ab*_ is the distance between vacated site *a* and a parent individual at site *b*. We use 1,000 sites on a line, with adjacent sites one unit distance apart (*d*_*a*,*a*+1_ = 1), leading to a fixed community size of 1,000 individuals. The reduced size relative to 21,000 individuals in the other models was used to keep this agent-based model computationally expedient. We explore a scenario with local dispersal (*d*_disp_ = 50), and one global dispersal (*d*_disp_ = 10^5^), i.e. where the probability of arrival is independent of distance to the parent.

### Competition-colonization tradeoff

Under this mechanism, changes to a population result from gains through recruitment and losses via death and displacement. Collecting the density-dependent and density-independent factors separately, one can show (see Supplementary Information) that the deterministic dynamics follow the equation

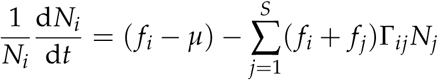

where *f*_*i*_ is the fecundity of species *i*, *µ* is a constant, and Γ_*ij*_ is the ability of a propagule of species *j* to displace an individual of species *i*.

The equation above is in Lotka-Volterra form. Therefore in parallel with our Lotka-Volterra scenarios, we place the density dependence on mortality. We thus set the probability that species *i* is chosen to die as Σ_*j*_ (*f*_*i*_ + *f*_*j*_)Γ_*ij*_ *N*_*i*_ *N*_*j*_/ Σ_*k*_ Σ_*l*_ (*f*_*k*_ + *f*_*l*_)Γ_*kl*_ *N*_*k*_ *N*_*l*_, while the probability it is chosen to recruit is *m f*_*i*_*q*_*i*_/ Σ_*j*_ *f*_*j*_*q*_*j*_ + (1 *- m*) (*f*_*i*_ *- µ*)*N*_*i*_/ Σ_*j*_ (*f*_*j*_ *- µ*)*N*_*j*_. Notice that high-fecundity species immigrate more frequently. The difference *f*_*i*_ - *µ* is an intrinsic growth rate, i.e. the net rate at which species grow in the absence of competition. Fecundities are log-uniform distributed between 1 and 6.2. We set *µ* = 1.

The tradeoff between fecundity and ability to displace is encoded in the displacement matrix Γ_*ij*_. If we arrange species in order of fecundity, Γ_*ij*_ will be approximately bottom-triangular, with Γ_*ij*_ ≈ 0 if *i < j* and Γ_*ij*_ ≈ 1 if *i > j*. That is, species with higher fecundity rarely displace those with lower fecundity, while low-fecundity species commonly displace high-fecundity ones. In the limit of an absolute tradeoff, lower-ranking species never displace and are always displaced by higher-ranking species. This limit has been shown to drastically inflate coexistence under this mechanism, despite the fact that such absolute tradeoffs are unlikely to occur in nature [74–76]. As such, here we use a probabilistic tradeoff: Γ_*ij*_ = 0.5 (1 *-* tanh[*s* (*f*_*j*_ *- f*_*i*_)]), which has a sigmoidal shape, is equal to 0.5 when *f*_*i*_ = *f*_*j*_, and asymptotes to 1 and 0 when *f*_*i*_ ≫ *f*_*j*_ and *f*_*i*_ ≪ *f*_*j*_, respectively. Parameter *s* controls the steepness of the transition, and hence the number of clusters. We set *s* = 0.15, which in the absence of immigration and stochasticity leads to c. 13 coexisting species (Fig S7).

### Clustering metrics: the gap statistic

We apply a variation of the gap statistic method [50], using two different clustering measures: k-means dispersion [48] and Ripley’s K function [77]. In general terms, the metric takes in the list of species traits and abundances, and returns the number of clusters (k-means) or average trait separation between them (Ripley’s K), as well as a z-score and a p-value. In Box 1 we summarize the k-means version and give a step-by-step recipe for its implementation, and illustrate it at work on two example communities. Both versions are described in detail in the Supplementary Information. The code for the k-means version is available on GitHub [78].

We assess statistical significance and degree of clustering by comparing a community against 100 null communities where we randomly shuffle local abundances across all species in the regional pool. This allows us to test specifically for a nonrandom association between traits and abundances, while keeping the observed abundance distribution fixed. One alternative null model is neutrality, whereby abundances follow a characteristic dispersal-limited multinomial distribution [79].

Because our models are stochastic, we run 100 replicates to account for variation within the same scenario. For each niche scenario we report the average z-score across 100 replicates, as well the percentage of replicates that were significantly clustered at level *p* < 0.05. To assess if a given run is significantly clustered we compared with the distribution across 100 nulls.

We check for false positives by testing our metric on neutral communities and communities where differences in species performance are due strictly to environmental filtering. From the former we expect significant clustering in circa 5% of runs when using a *p* = 0.05 cutoff, and from the latter we expect that species will cluster around the favored trait. Thus we distinguish niche differentiation from neutrality and pure environmental filtering by the presence of multiple clusters as opposed to none or a single one.

### Box 1: Detecting clusters with k-means and the gap statistic

Here we describe our metric based on coupling the k-means clustering algorithm [48] with the gap statistic method [50]. Conceptually, we assess the degree of clustering by measuring the total trait distance between species in the same cluster, summed over all clusters. Smaller values indicate tighter clustering. However, we cannot use this index directly unless we know the number of clusters to look for, as the index naturally declines with the number of clusters considered. To circumvent this, we compare this total distance to that found in null communities, and find the number of clusters that maximizes the difference from the null. This maximal difference is the statistic.

Stepwise, we proceed as follows. We start with a candidate number of clusters, *k*. Out of all possible ways to assign the observed species to *k* clusters, the k-means algorithm finds the one that minimizes within-cluster trait dispersion: *D*_*k*_ = Σ_*C*_ Σ_*i*,*j∈C*_ *n*_*i*_*n*_*j*_*d*_*ij*_, where C refers to a cluster (1 ≤ *C* ≤ *k*), *n*_*i*_ is the abundance of species *i*, *d*_*ij*_ is the trait distance between species *i* and *j*. We define the clustering index *F*_*k*_ = *-* log(*D*_*k*_), such that *F*_*k*_ is high for low dispersion *D*_*k*_. We then repeat this step across a set of 5,000 null communities, and the difference in *F*_*k*_ between the observed community and the mean null value is the gap for *k* clusters, 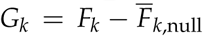. We then test a range of numbers of clusters and find the one that maximizes the gap. This maximal value (the peak in the gap curve in Fig B1B) is the gap statistic, *G* = max(*G*_*k*_), and the value of *k* at which it occurs is the estimated number of clusters *K*. Note that the gap step is needed because *F*_*k*_ will always increase with *k*: as we increase the number of clusters, the average cluster becomes smaller and therefore within-cluster dispersion decreases. By comparing against null communities, the gap method selects the number of clusters that most increases the clustering index beyond those expectations.

We then perform the same routine on each of the null communities to obtain a null distribution of gap statistics, from which we extract significance (p-value) and standardized effect size (z-score). The z-score is *Z* = (*G - µ*)/*σ*, where 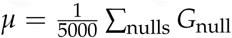 is the mean of the null gap statistics and 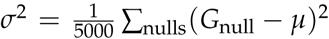 is the variance. The p-value is the proportion of null communities with a higher gap statistic than the observed community, 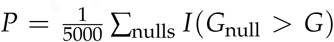, where the indicator function *I* is 1 if its argument is true, and zero otherwise.

Fig B1 shows two test communities, and the respective results from our metric. The first community (Fig B1A) is constructed to contain four clusters, whereas in the second community (Fig B1C) abundances are unrelated to traits. The metric correctly diagnoses both (compare Fig B1B, D).

**Figure B1:**
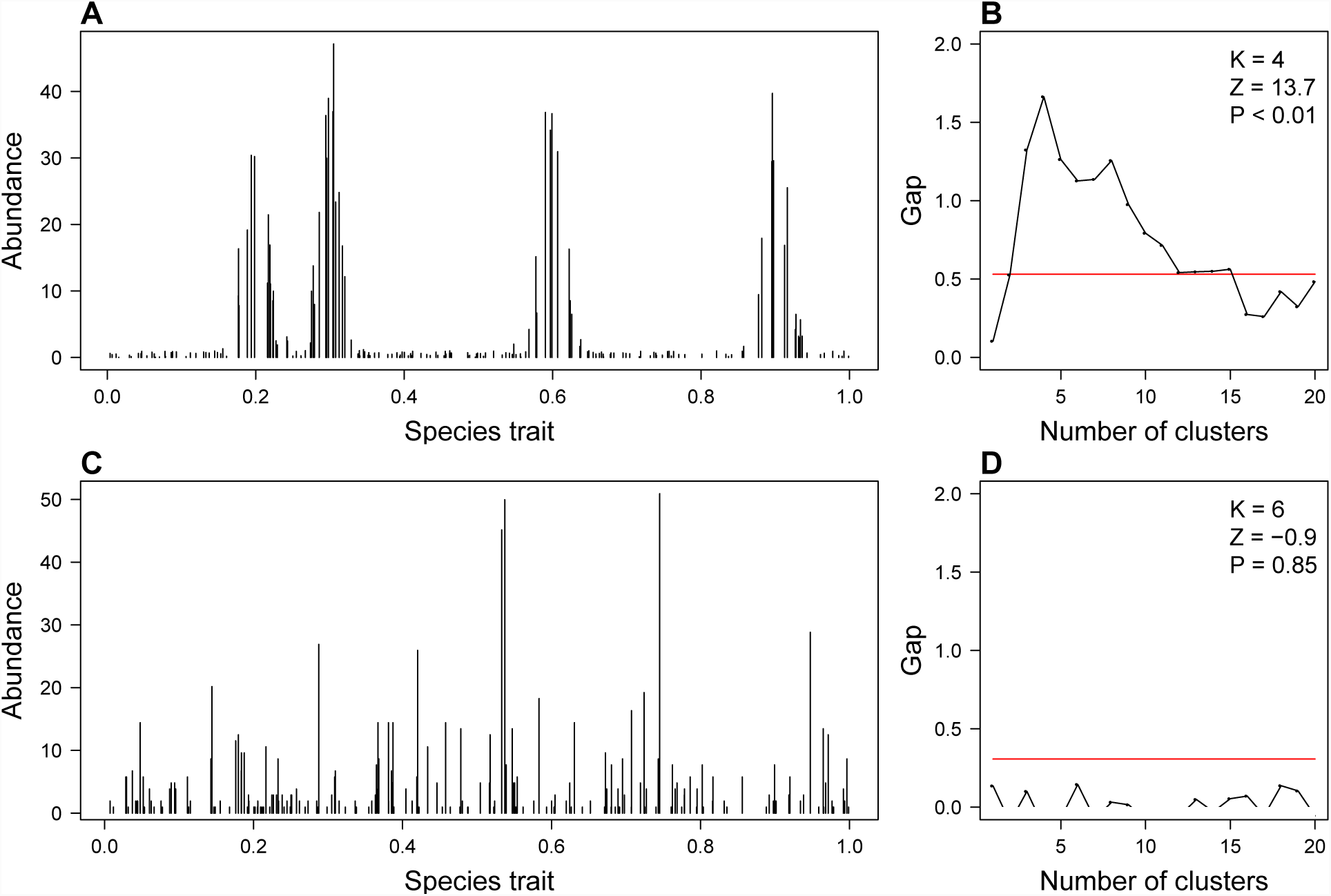
**A**: Hypothetical species assemblage representing a community under niche differentiation + immigration, with visible clustering. **B**: Plotting the dispersion gap against the number of clusters, we see a clear peak at *K* = 4 clusters. The gap value at the peak, *G*_4_ = 1.66, is the gap statistic. It far exceeds the 95th percentile of the gap statistic across the null communities (red line), leading to a very high z-score and a very low p-value. **C**: Neutral community with same size as A, but abundances unrelated to traits. **D**: Corresponding gap curve indicates that no number of clusters gives a gap beyond null expectations. Witness the low z-score and high p-value. [*Numerical details*: Community A: 200 species traits were uniformly drawn between 0 and 1. We then assigned a species to each of 1,000 individuals, with probability of choosing species *i* given by 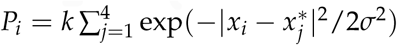, where *x*_*i*_ is the trait of species *i*, **x**^*∗*^ = {0.2, 0.3, 0.6, 0.9} are the centers of the four clusters, *σ* = 0.02 is the cluster width, and *k* is a normalizing constant ensuring 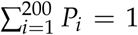. Finally, abundances are boosted by a random small amount to represent immigration. Community C: we drew species abundances from a log-series distribution with parameters *N* = 1, 000 and *α* = 50, and assigned them to randomly chosen trait values.]

### Glossary

#### Cluster

a set of species with similar trait values and relatively high abundance, separated from other such sets by sparsely populated regions in trait space.

#### Competition-colonization tradeoff

a niche mechanisms commonly invoked for plant coexistence. Species may offset disadvantages in one-on-one contests for recruitment via superior ability to colonize freshly available habitat, or vice-versa [46].

#### Ecological drift

ecological analogue to genetic drift, whereby changes in the abundance of a species occur via random sampling from a finite population, independent of fitness or competitive standing related to species traits [38].

#### Limiting similarity

a classical theoretical concept that has taken on multiple meanings over time [5,80], here interpreted as greater-than-chance trait differences between adjacent species on the niche or trait axis. While competing species were traditionally expected to display limiting similarity, the demonstrated robustness of clusters to demographic stochasticity, their persistence under immigration, and their appearance under disparate niche mechanisms suggest that clusters may be more common than limiting similarity.

#### Mass effects

phenomenon in source-sink dynamics whereby a locally disfavored population can be sustained via immigration from populations favored elsewhere [81].

#### Niche, niche strategy, niche differences

a concept in community ecology pertaining to idiosyncrasies in how a species responds to and in turn affects the environment, resources, and/or other species [2]. Traditionally used to describe species differences in environmental preferences or in strategies for procuring resources, avoiding enemies, etc. In modern co-existence theory, “niche differences” are operationally defined as species differences that stabilize competitive coexistence, which requires stronger competition within than between species[6,7]. Stable competitive coexistence occurs when each species can grow from low abundance in the presence of the others.

#### Niche axis

the axis of interspecific variation pertaining to niche strategies and niche differences. For example if insectivorous bird species differ in their preference for prey of different sizes, preferred prey size would form a niche axis. In contrast, in the competition-colonization tradeoff, the axis runs from competition specialist (high-quality low-number seeds) to colonization specialist (low-quality high-number seeds).

#### Niche mechanism

any set of circumstances that stabilizes coexistence, typically by ensuring that competition is stronger within species than between them [7]. In a niche mechanism, the nature of species interactions, plus sometimes the presence of tradeoffs, lead to opportunities for species to differ in their interaction with limiting factors [82]. Examples include specialization to different prey and/or different abiotic environments.

#### Optimal niche strategy

In a closed community, when the current number of species exceeds the number of species that can stably coexist, only those with optimal niche strategies will persist indefinitely. Optimal niche strategies are sufficiently separated from one another on the trait axis to enable stable coexistence. The particular set of optimal niche strategies that emerges from the competitive process can be determined in part by external circumstances such as which resources are in greater supply, or simply by the starting conditions or order of arrival of species to the community. Mathematically, models of competition among species on niche axes typically have a pattern-forming instability [52], whereby species with optimal strategies become increasingly abundant while all others dwindle towards exclusion.

#### Stabilization vs. equalization

a dichotomy in modern coexistence theory [7] whereby competitive exclusion can only be mitigated via mechanisms that either stabilize the community (niche mechanisms), or reduce average fitness differences between species (equalizing mechanisms). While the concept is a useful theoretical construct, it can be difficult or impossible to calculate the degree of stabilization and equalization in multispecies communities, or to assign trait differences to one or the other [62] (but see [83] for an update to the theory).

## Supporting information

## Acknowledgments

We thank David Ackerly for helpful discussions, and James O’Dwyer, Brian Sedio, Stephen Smith, and Lacey Knowles for comments on previous versions of the manuscript. This work was supported by the National Science Foundation under grant 1038678, “Niche versus neutral structure in populations and communities” funded by the Advancing Theory in Biology program, as well as by the Danish National Research Foundation through support of the Center of Macroecology, Evolution and Climate (grant DNRF96), and by the Miller Institute for Basic Research in Science at the University of California, Berkeley.

## Footnotes

1 In fact the model can be cast in Lotka-Volterra form, and one can show that the net competitive impact, while asymmetric, is stronger between more similar species (Fig S7).

