## Supplementary material for "Generalizing clusters of similar species as a signature of coexistence under competition"

### Supplementary Information

#### Supplementary Figures

Figure S1: **Community under environmental filters but no niche mechanism shows a single cluster.** **A**: Example simulation outcome of Lotka-Volterra stochastic dynamics with neutral competition coefficients, $\alpha_{ij} = 1$, such that there is no niche mechanism, and intrinsic growth rates given by $r_i = x_i(1 - x_i)$, where $x_i$ is the trait of species $i$. The latter represents environmental filtering for species with intermediate traits. **B**: Corresponding gap curve, showing gap index for each number of clusters between 1 and 20, has a clear maximum at 1 cluster. The estimated number of clusters is therefore $K = 1$. The gap statistic is well above the 95% quantile of the null distribution (red line), indicating significance at $P < 0.05$. All 100 replicates of this scenario were clustered, with a single cluster detected by the k-means metric in all cases.

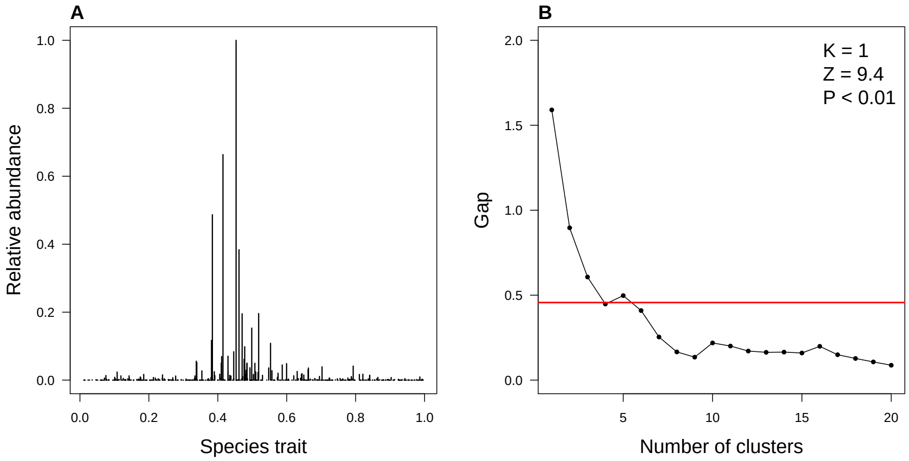

Figure S2: **Lotka-Volterra niche scenarios with environmental filtering**. **A**: We implemented environmental filters via a modal relationship between intrinsic growth rate $r$ and species traits $x_i$ as follows: $r = 0.5$ (no filtering, black); $r_i = x_i(1 - x_i)$ (red); $r_i = \exp(-(x_i - 0.5)^2/\sigma^2)$ with $\sigma = 0.5$ (green), 0.2 (blue), 0.1 (magenta). **B**: Under no filtering, the community shows no overarching abundance trend. **C-F**: Under increasing filter intensity, communities show increasingly steeper abundance trends. Our metrics detected multiple clusters in 10/10 replicates of B-D, but only 2/10 in E and 1/10 in F, with the remaining replicates having a single cluster. When no niche mechanism is at play, such that species compete neutrally but still differ by intrinsic growth rates, all replicates result in a single cluster centered on the species with the highest intrinsic growth rate (Fig S1).

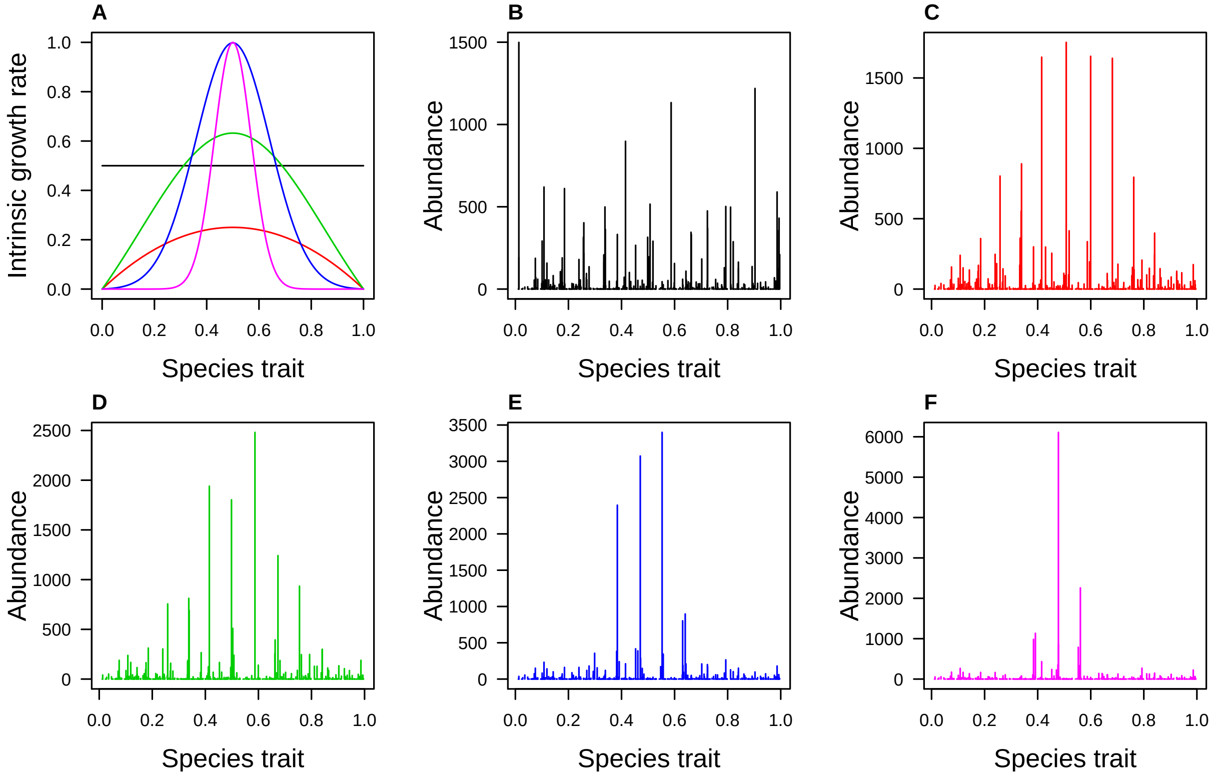

Figure S3: **Effects of immigration rates and regional diversity.** Example Lokta-Volterra communities with increasing immigration rate $m$ (left) and regional diversity parameter $\theta$ (right).

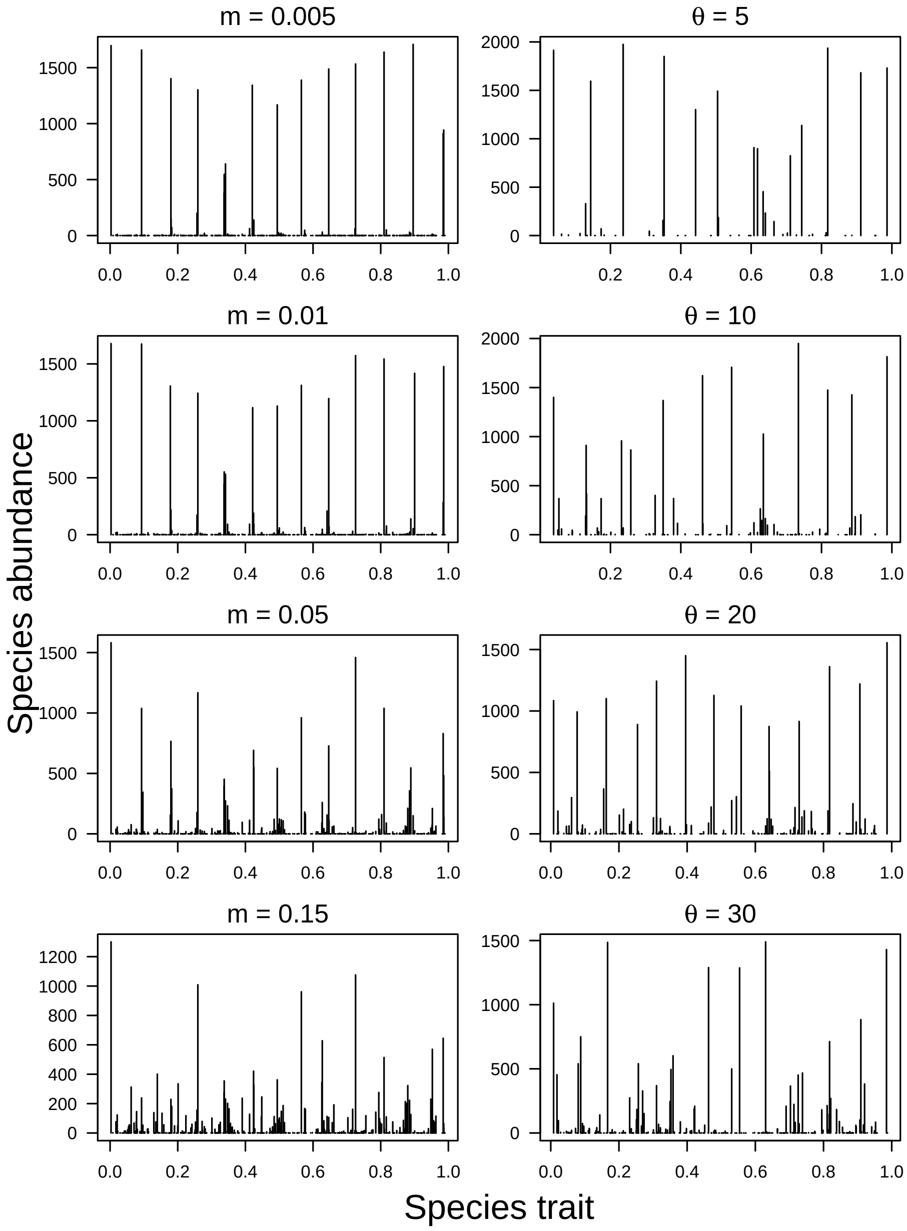

Figure S4: **Lotka-Volterra and other niche communities under lower immigration**, $m = 0.01$. Compared with $m = 0.08$ (Fig 2), communities show a similar number of clusters but fewer species per cluster, and gaps between clusters are more rarified.

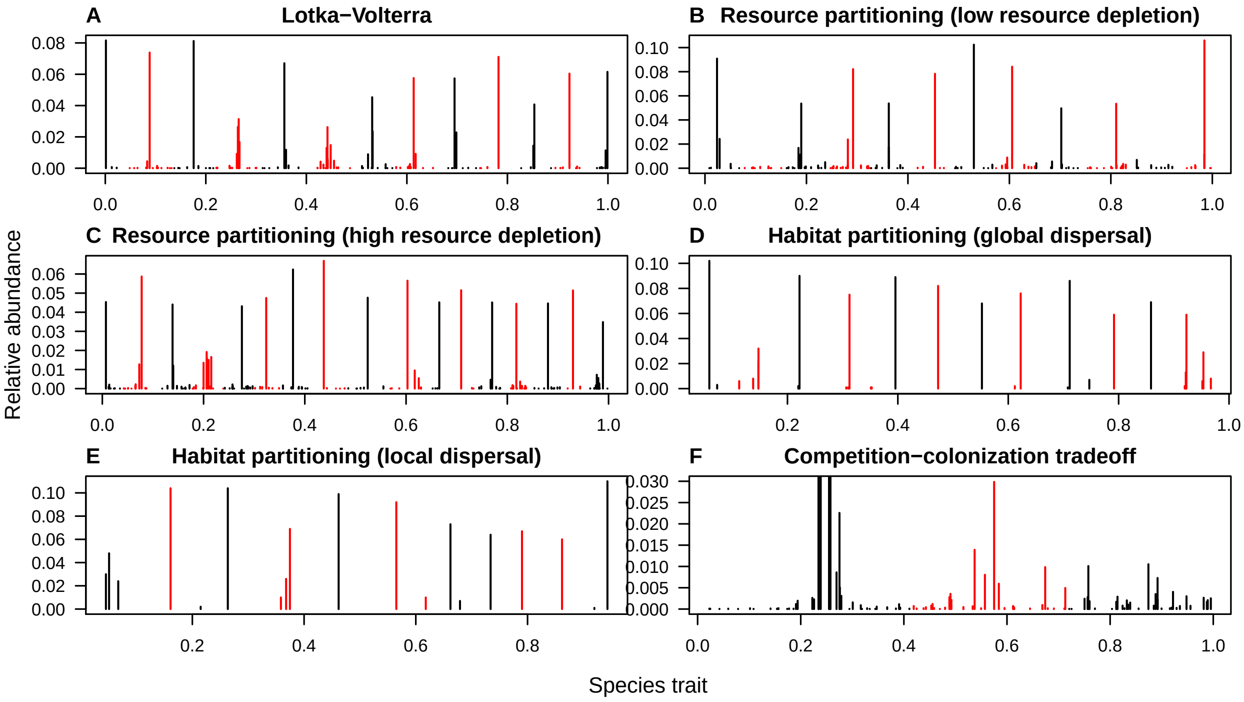

Figure S5: **Clustering results under lower immigration**, $m = 0.01$. Results are similar to $m = 0.08$ (compare with Fig 3), although z-scores and significance are often higher, particularly for resource-partitioning communities, and habitat-partitioning communities under global dispersal. This indicates that higher immigration is drowning the pattern in these niche scenarios.

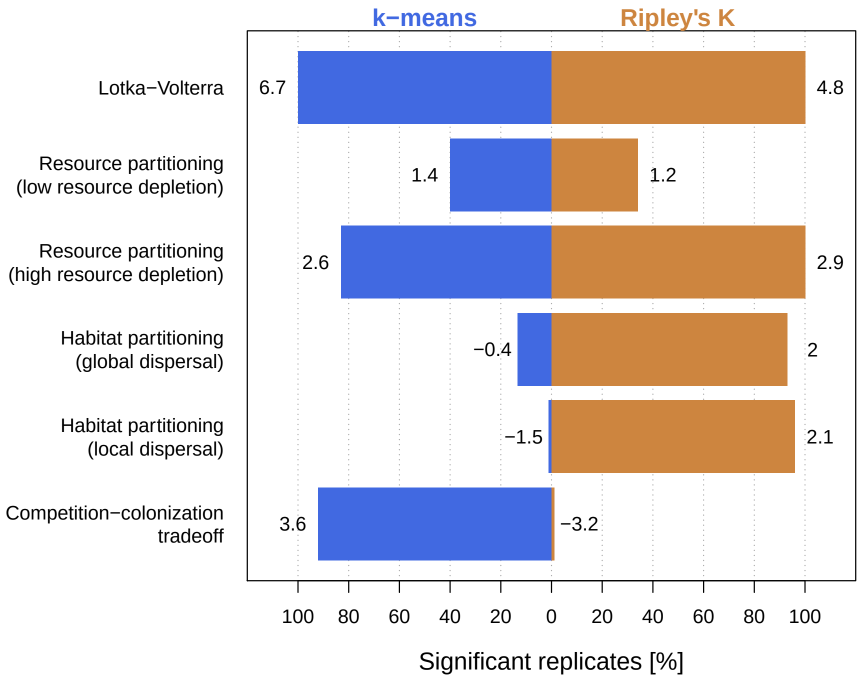

Figure S6: **Effects of resource depletion.** Example communities where consumers partition resources, under low and high resource depletion (left and right columns, respectively). Resources are shown on top, consumers at the bottom. Under high resource depletion, gaps left by resource extirpation cause corresponding gaps among consumer species. These gaps strengthen the clustering pattern relative to the low depletion scenario. ($m = 0.08$, c. 400 regional species.)

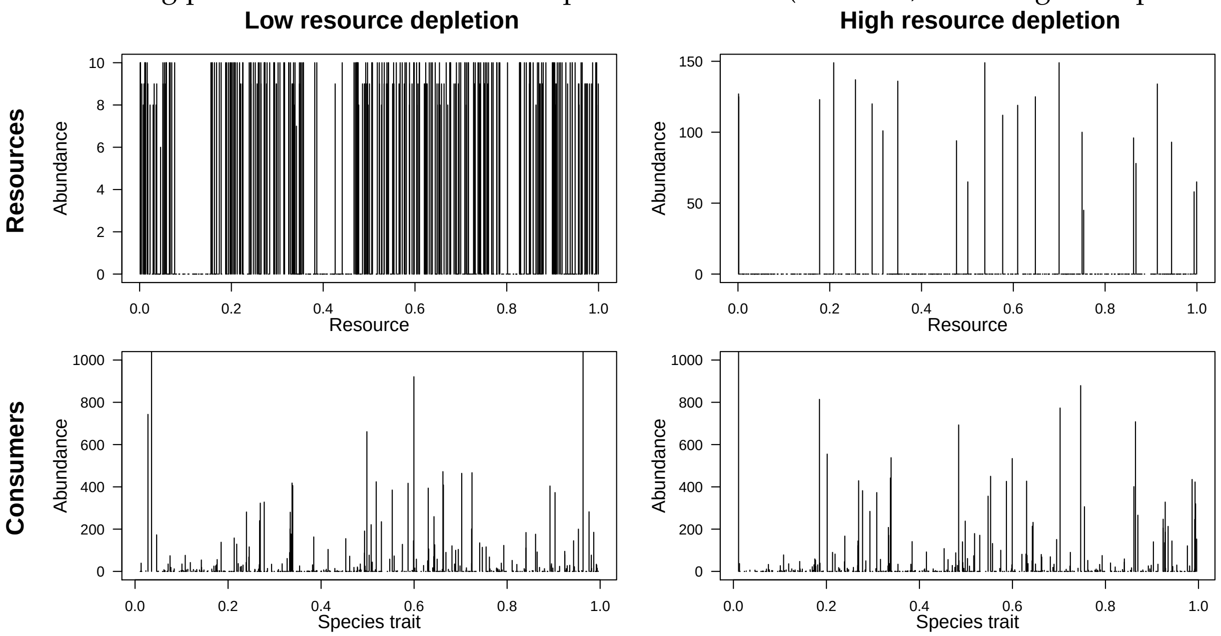

Figure S7: **Competition-colonization tradeoff, deterministic implementation without immigration.** **A**. Transient state shows visible clustering. **B**. Equilibrium state, showing abundances of the coexisting species, which are those that dominate their respective clusters in the transient state (marked with red dots in A). Legend shows maximum eigenvalue of the Jacobian of the equilibrium, indicating dynamical stability of the equilibrium. **C**. Competition kernel shows the strongest competitors on focal species, whose traits are shown in legend, are species with similar traits, despite the competitive hierarchy. This contributes to stabilization of the community. **D**. For each species (traits plotted on x-axis), the position of the peak of the competition kernel (i.e. the species with the strongest net competitive impact on it) is plotted on the y-axis. The proximity of the curve to the one-to-one line (dotted line) throughout the trait axis shows that the competition-colonization tradeoff stabilizes the community, thus acting as a niche mechanism. The kernel maxima plotted here also explain the absence of coexisting species below trait $\simeq 0.2$, as the strongest competitors of those species have higher fecundity than themselves (solid curve is above dotted line), thus being both more competitive and more fecund. The wide gap between the first cluster and the other clusters is also reflected in the relatively larger distance from the one-to-one curve in that region of the trait axis.

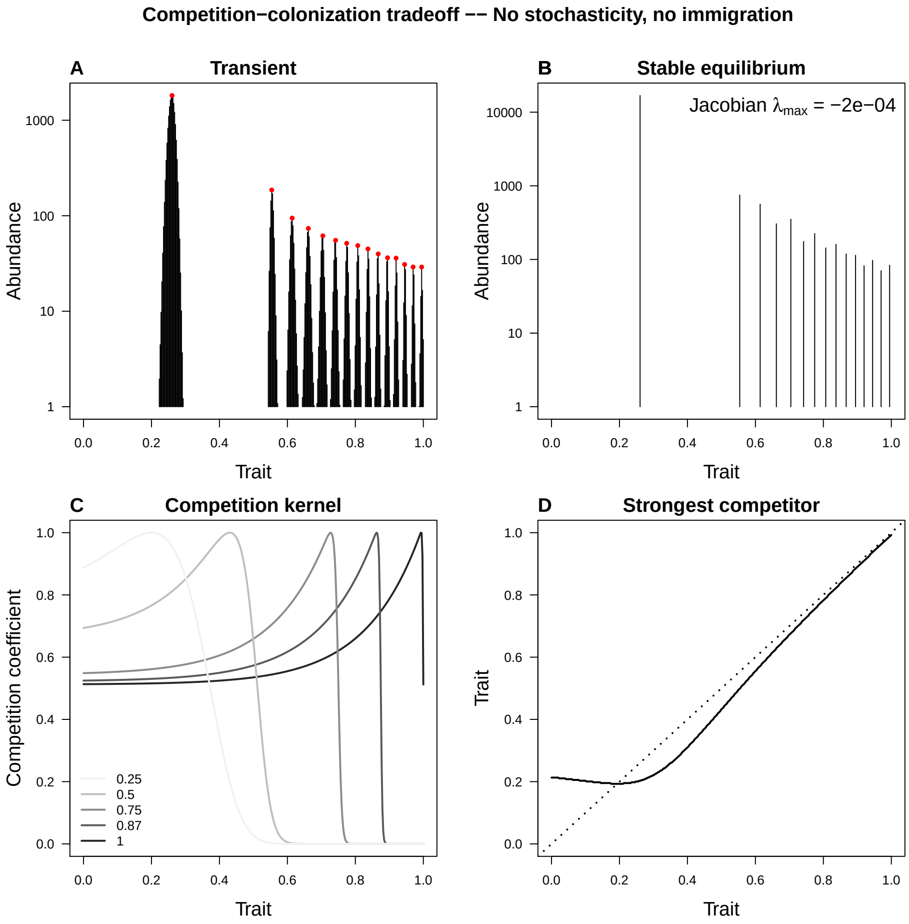

#### Additional Methods

##### Competition-colonization tradeoff model

This model was originally formulated as follows [84]

$$\frac{1}{p_i}\frac{\mathrm{d}p_i}{\mathrm{d}t} = f_i(1 - \sum_{j=1}^{i} p_j) - \sum_{j=1}^{i-1} f_j p_j - \mu, \tag{S1}$$

where $p_i$ is species $i$'s relative abundance, and $f_i$ is its fecundity (i.e. number of propagules per individual per unit time). The first term represents recruitment in available sites—i.e. all sites not currently occupied by species $i$ or stronger competitors; the second term represents displacement by stronger competitors; the last term is intrinsic mortality $\mu$, here assumed identical for all species.

We can gather the density-independent and the density-dependent terms separately:

$$\frac{1}{p_i}\frac{\mathrm{d}p_i}{\mathrm{d}t} = (f_i - \mu) - \sum_{j=1}^{S} \Theta_{ij}(f_i + f_j)p_j, \tag{S2}$$

The step function $\Theta_{ij}$ (equal to 1 if $i > j$, 0 if $i < j$, and 0.5 if $i = j$) encodes the strict competitive hierarchy. It has been shown [75] that the strict hierarchy is unrealistic (because arbitrarily similar species will have large differences in competitive ability) and drastically inflates coexistence. Here we use instead a probabilistic, gradual hierarchy: the higher the difference in rank, the higher the likelihood of displacement by the better competitor. We do this by replacing $\Theta_{ij}$ with the continuous function $\Gamma_{ij} = 0.5\,(1 - \tanh[s(f_j - f_i)])$, which is equal to 0.5 when $f_i = f_j$ and asymptotes to 1 and 0 when $f_i \gg f_j$ and $f_i \ll f_j$, respectively. The parameter $s$ controls the steepness of the hierarchy, and hence the degree of coexistence. (We recover the step function when $s \to \infty$.)

Notice that Equation (S2) is in the Lotka-Volterra shape, $\frac{1}{N_i}\frac{\mathrm{d}N_i}{\mathrm{d}t} = r_i - \sum_j A_{ij}N_j$, where the effective intrinsic growth rate $r_i = b_i - d_i$ is the difference between intrinsic fecundity and mortality. We use that correspondence in our stochastic formulation, where we place the density dependence in the deaths.

##### Detailed description of clustering metrics

Our method proceeds in the following general steps:

1. Given the data and a certain parameter value $\varphi$ within a provided range $[\varphi_{\min}, \varphi_{\max}]$, calculate a clustering measure $F(\varphi)$.

2. Repeat step 1 in each of $n$ null communities, $\widetilde{F}_i(\varphi)$, $i = 1, 2, \ldots, n$, and take the mean, $\overline{\widetilde{F}}(\varphi) = \frac{1}{n}\sum_i \widetilde{F}_i(\varphi)$.

3. The difference $G(\varphi) = F(\varphi) - \overline{\widetilde{F}}(\varphi)$ is the *gap* at that parameter value.

4. Repeat steps 1-3 for all parameter values within $[\varphi_{\min}, \varphi_{\max}]$.

5. The gap statistic is the extremum (maximum or minimum) of the gap function, $G = \underset{\varphi}{\text{extr}}\, G(\varphi)$.

6. Obtain a z-score and a p-value by repeating steps 1-5 on each of the null communities, which provides a null distribution for $G$. The z-score is then $Z = (G - \mu_{\widetilde{G}}) \,/\, \sigma_{\widetilde{G}}$ and the p-value is $P = \frac{1}{n}\sum_i I(\widetilde{G}_i \geq G)$, where $\widetilde{G}_i$ is the gap statistic obtained for the $i$-th null assemblage, $\mu_{\widetilde{G}}$ and $\sigma_{\widetilde{G}}$ are the mean and standard deviation of those values, and index function $I$ is 1 if its argument is true and 0 otherwise.

Note that this recipe works with different clustering measures, $F$. Here we use two: k-means dispersion [48] and Ripley's K function [77]. In the k-means version, the parameter $\varphi$ is the number of clusters, and the value $\hat{\varphi}$ at which the gap is maximal is the estimated number of clusters in the community. In the Ripley's K version, $\varphi$ is the trait distance between pairs of individuals, and the value at which the gap is minimal is the average distance between clusters.

Note also that we can use any number of traits to describe our species, as long as we can define a "distance" between species (e.g. Euclidean distance in a high-dimensional trait space, or simple trait differences on a single trait axis).

##### Gap statistic via k-means

Here the parameter $\varphi$ is the candidate number of clusters $k$, and we use the k-means clustering algorithm [48]. For a given $k$, the algorithm finds the partition of individuals into $k$ groups that minimizes the within-group dispersion. Let $D_k$ be the total pairwise squared distance between members of each group, i.e., $D_k = \sum_{C=1}^{k}\sum_{i,j\in C} n_i n_j d_{ij}^2$, where C refers to a cluster, $n_i$ is the abundance of species $i$, $d_{ij}$ is the trait distance between species $i$ and $j$. The k-means algorithm finds the species-to-cluster assignment[1] that minimizes $D_k$. It starts with randomly chosen trait values in the local community as possible cluster centers, then puts species into the cluster whose center is the closest to them, then recalculates cluster centers, and so on until the algorithm converges or subsequent changes in $D_k$ fall below a specified threshold. Since the result can depend on the starting point, we carry out this procedure from a variety of randomly chosen cluster centers, and take the final cluster arrangement with the lowest $D_k$ across different starting points[2]. We then set $F(\varphi) = F_k = \log(1/D_k)$ as the clustering measure to be maximized with the Gap statistic[3]. See Box 1 for a step-by-step recipe for this metric.

[1]In our model, all conspecific individuals have the same trait value, and therefore necessarily belong in the same cluster. For efficiency we modified the k-means algorithm to arrange all individuals of a species together.

[2]If a cluster ended up empty in this approach, the arrangement was not included in calculating the minimum $D_k$. We assessed the number of starting points needed by verifying that the resulting $D_k$ changed little if more starting points were added. This number was relatively similar across model scenarios. We used a single starting point for 1 cluster (for which the arrangement is independent of the number of starting points), 1,000 starting points for 2-5 clusters, 5,000 starting points for 6-10 clusters, 10,000 starting points for 11-15 clusters, and 100,000 starting points for 16-20 clusters. The exception was the habitat partitioning model, where empty clusters occurred frequently, and hence we used fewer starting points to maintain reasonable computational time.

[3]Note that it does not make sense to compare $F_k$ directly across different $k$'s because $F_k$ necessarily increases with $k$, as the average within-cluster distance is always lower for higher numbers of clusters. By comparing against null assemblages, the gap method finds the biggest increase in goodness of fit *beyond* what is expected from the increase in $k$. The reason we use $\log(1/D_k)$ rather than simply $1/D_k$ is that the expected increments in $1/D_k$ with increasing $k$ are multiplicative rather than additive (see [50] for more details).

**Gap statistic via Ripley's K**

The approach with Ripley's K is to count the number of pairs of individuals within each distance in trait space and determine whether or not it deviates significantly from what we would expect if the same set of abundances were randomly distributed among the present species. In particular, we look for significant open spaces between clusters, i.e. distances at which Ripley's K is surprisingly low. The parameter $\varphi$ here is therefore the trait distance $d$.

Ripley's K function at distance $d$ is defined as $K(d) = \frac{\sum_{i \neq j} I(d_{ij} < d) N_i N_j}{\sum_{i \neq j} N_i N_j}$, where $N_i$ is the abundance of species $i$, $d_{ij}$ is the distance in trait space between species $i$ and $j$, and function $I(.)$ is the indicator function, equal to 1 if its argument is true and 0 otherwise. We then calculate Ripley's K in each of our null assemblages, and define the Ripley's K clustering measure as the standardized K function: $F(d) = K(d)/\sigma_{\widetilde{K}(d)}$, where $\sigma_{\widetilde{K}(d)}$ is the standard deviation of the null values. The standardization compensates for the natural increase in the variance of $K(d)$ at large $d$ (see Fig S8). The gap function is then $G(d) = K(d)/\sigma_{\widetilde{K}(d)} - \mu_{\widetilde{K}(d)}/\sigma_{\widetilde{K}(d)}$.

Because we are interested in significantly low counts, we define our Ripley gap statistic as the minimum value of $G(d)$ across candidate distances, $G = \min_d G(d)$. This is analogous to our implementation with the k-means algorithm (see Table S1 for a direct comparison between the two implementations).

Fig S8 shows the metric at work on an example of a Lotka-Volterra community, a community under neutral competition and environmental filtering, and a purely neutral community. When the community has significant open spaces (Fig S8A), i.e. a lower $G$ than the null communities, it suggests multiple clusters in trait space of high abundance species, with a dearth of species in between them. This is what we would expect from a community shaped by niche differentiation. When instead it is significantly lacking in gaps (Fig S8B), that indicates a single clump of abundant species, as could be expected when the environment filters for a single trait value.

Note that while k-means and Ripley's K can characterize a community in terms of clustering structure, the gap statistic tells us whether that structure is significant compared to a null model. Furthermore, while clustering measures will typically depend on a parameter (number of clusters in the case of k-means, distance between clusters in the case of Ripley's K), the gap statistic removes the parameter by comparing the data to the null model across the parameter's range.

Table S1: **Comparison of our two implementations of the gap method**. Both apply the gap method using different measures of dispersion. $W(k)$ is the k-means dispersion for $k$ clusters; $K(d)$ is Ripley's K at trait distance $d$; $F(k)$ ($F(d)$) is the dispersion for $k$ clusters (distance $d$); $I$ is the indicator function, equal to 1 if its argument is true and zero otherwise; $N_i$ is the abundance of species $i$; $G(k)$ ($G(d)$) is the gap function for $k$ clusters (distance $d$); $\mu_{\widetilde{X}}$ and $\sigma_{\widetilde{X}}$ are the mean and standard deviation of quantity $X$ taken across the null communities; $\hat{d}$ is the distance at which $G(d) = G$. $n$ is the number of null communities.

| | **k-means** | **Ripley's K** |
|---|---|---|
| Clustering measure | $F_k = \log(1/W(k))$ | $F(d) = K(d)/\widetilde{\sigma}_K(d)$ |
| Gap function | $G_k = F_k - \mu_{\widetilde{F}_k}$ | $G(d) = F(d) - \mu_{\widetilde{F}(d)}$ |
| Gap statistic | $G = \max_k G_k$ | $G = \min_d G(d)$ |
| Estimated number of clusters | $k$ such that $G_k = G$ | Number of abundance-peaks within distance $\hat{d}$ of each other |
| Z-score | $Z = (G - \mu_{\widetilde{G}}) \,/\, \sigma_{\widetilde{G}}$ | $Z = (G - \mu_{\widetilde{G}}) \,/\, \sigma_{\widetilde{G}}$ |
| P-value | $P = \frac{1}{n}\sum_i I(\widetilde{G}_i \geq G)$ | $P = \frac{1}{n}\sum_i I(\widetilde{G}_i \geq G)$ |

Figure S8: **Top**: Abundances plotted against trait values in a sample replicate of (**A**) the Lotka-Volterra niche model, (**B**) environmental filtering without a niche mechanism, and (**C**) the neutral model. **A1-C1**: Ripley's K vs. trait distance in the same replicates. Values for the observed community are plotted in black. The %95 confidence interval of $K(d)$ among the null communities (obtained by reshuffling abundances across species) is shaded in gray. **A2-C2**: same results with mean null value at each distance subtracted out, $K(d) - \widetilde{\mu}_K(d)$. **A3-C3**: rescaled by the variance at each distance $G(d) = K(d)/\widetilde{\sigma}_K(d) - \widetilde{\mu}_K(d)/\widetilde{\sigma}_K(d)$. The red line is the threshold for a significantly low gap statistic ($p < 0.05$), and the triangle indicates the distance with the most surprisingly low density of pairs, i.e. the distance $d$ where the Ripley gap statistic $G$ is achieved, $G(d) = G$.

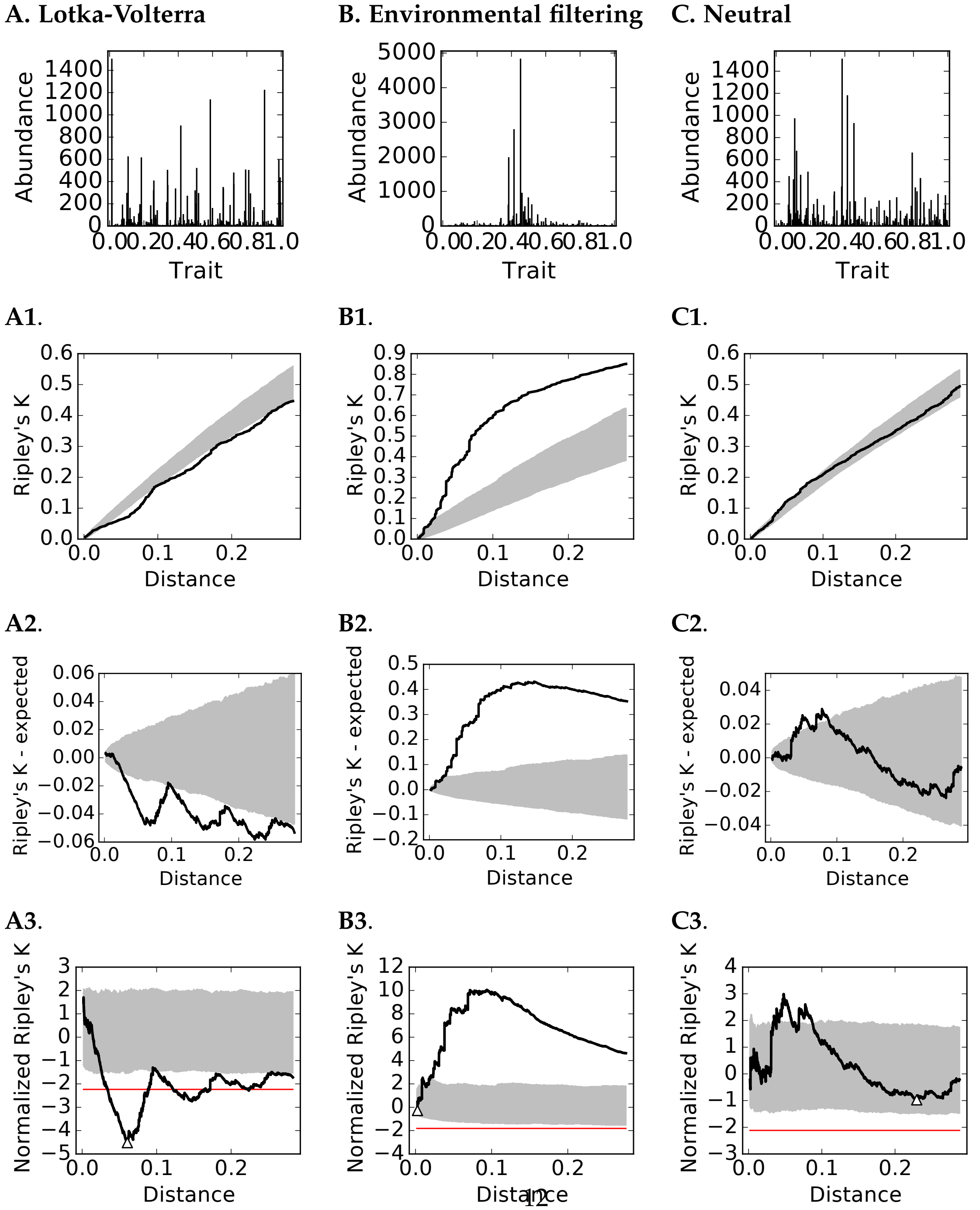
